# Altered Orbitofrontal Cortex Activation and Gaze Patterns to Happy Faces in Autistic Children Predict Adaptive Difficulties but Challenge the Social Motivation Hypothesis

**DOI:** 10.1101/2023.12.04.569509

**Authors:** Mengyuan Yang, Lan Zhang, Zijie Wei, Pingping Zhang, Lei Xu, Lihui Huang, Hong Li, Yi Lei, Keith M. Kendrick, Juan Kou

## Abstract

Individuals diagnosed with Autism Spectrum Disorder (ASD) generally have altered responses to social reward, which the “social motivation” hypothesis posits as the primary contributor to their social deficits. Aberrant perception of rewarding faces is considered to be one of the most significant markers of this hypothesis. The orbitofrontal cortex (OFC) plays an important role in modulating reward and arousal responses to happy faces but few studies have investigated whether young autistic children with limited cognitive or language ability have altered neural and behavioral responses to reward related tasks. We therefore conducted an eye-tracking task where autistic (n = 36) and typically developing (n = 36) young children (aged 2.5-6 years) viewed the faces of happy children or of simplified faces (emoticons), combined with OFC activation measurement using functional near infrared spectroscopy (fNIRS), to compare ASD and typically developing (TD) children. Results revealed no differences between the two groups for time spent viewing happy faces although TD children spent more time looking at the eyes of real faces and at the mouths of emoticons whereas children with ASD did not. Children with ASD showed a greater pupil diameter (PD) and OFC activation compared to TD children during presentation of happy faces. Indeed, greater PD and OFC responses were predictive of greater adaptive behavior difficulties and severity of symptoms. Our results therefore demonstrate different gaze patterns and greater arousal and brain reward system responses to happy faces in young children with ASD which are incongruent with social motivation hypothesis.

**Availability of data:** Our ethics approval does not permit public archiving of individual anonymized raw data. Those who wish to access the raw data should contact the corresponding author. Access will be granted to named individuals who adhere to the ethical procedures governing the reuse of sensitive data. They must complete a formal data sharing agreement to obtain the data. The data that support the main group different finding figures related anonymized data of this study are openly available via the Open Science Framework (OSF) Repository https://osf.io/sxkhc/?view_only=6eb51e64a9b54fce909b5ff6eacdc668

**Ethics approval statement:** The study was conducted in accordance with the principles of the Declaration of Helsinki. It was approved by the Local Ethics Committee (number 202165).

**Research Highlights:**

● The present study examined whether altered social reward processing in ASD is caused by reduced behavioral and neural responses to rewarding stimuli.
● Typically developing children showed a preference for viewing specific informative facial features (such as child eyes or emoticon mouths), children with autism did not.
● Children with autism enhanced arousal (pupil dilation) and brain reward (orbitofrontal cortex) responses to happy child faces.
● Altered, gaze patterns, arousal and brain reward responses were highly predictive of adaptive behavior abilities and autistic symptom severity.

## 1 Introduction

Autism Spectrum Disorder (ASD) is a neurodevelopmental condition, identified by difficulties in social communication and repetitive and restrictive behaviors or interests (American Psychiatric Association, 2013). A large number of studies have reported that individuals with autism show a decreased attention to and preference for social stimuli when compared to those with typical development (Dubey et al., 2015; Chita-Tegmark, 2016; Clements et al., 2018). Individuals with ASD often appear to have a lack of social motivation while processing social information which has led to a hypothesis that symptoms may be particularly contributed to by this reduced social motivation (Chevallier et al., 2012; Kohls, et al., 2014). Perception of social reward (eg: happy faces), one of the richest and most powerful factors involved in conveying social information during social interactions and its alteration in autism has been considered to be one of the most significant markers supporting this hypothesis (Jack & Schyns, 2015; Frazier et al., 2017; Chevallier et al., 2012).

### 1.1 Altered gaze patterns during processing of faces in ASD

Individuals with ASD show less attention to whole faces and notably eye regions that are crucial for accurate social perception regardless of age (4 months to 40 years old)(Frazier et al., 2017). A ground-breaking eye-tracking study determined that adolescents and adults with ASD had a tendency to look more towards the mouth and less towards the eyes compared to TD individuals (Klin et al., 2002). This study drew a lot of attention, resulting in the so-called “excess mouth/diminished eye gaze hypothesis” in subsequent ASD research (Guillon et al., 2014; Hutchins & Brien, 2016; Pierce et al., 2001; Nomi & Uddin, 2015).However, some studies have reported that ASD and TD individuals both spend more time looking at the eyes than the mouth without showing any group differences in either time spent viewing these features or in the ratio of attention towards them (Vettori et al., 2020; Rutherford et al., 2008). A recent eye-tracking study was conducted on a large sample of toddlers with and without ASD (N = 385, 11-47 months old) and found similar patterns of eye and mouth within the context of viewing child-friendly vignettes (Kwon et al., 2019). Several factors have been suggested to account for disparities between the findings of different studies, including the degree of symptom severity, differences in outcome indices as well as differences in age and cognitive competence.

### 1.2 Altered social motivation in ASD

An influential preliminary study reported that social stimuli were perceived as having a low salience and reward value in individuals with ASD leading to hypo-responsiveness to social stimuli and difficulties in social information processing (Chevallier et al., 2012). It has been suggested that altered reward circuitry in the brains of autistic children results in them not experiencing social stimuli in the same way as neurotypical children, especially in the context of social reward (Scott et al., 2010; Kohls et al.,2018);. A lack of social motivation may therefore lead to reduced social learning experiences, which could impede the development of language and social skills.

Extensive research has been conducted to identify the neural mechanisms associated with ASD, with a particular focus on the orbitofrontal-amygdala circuit and the locus coeruleus-norepinephrine (LC-NE) system, both of which are believed to be important for etiology and treatment (Bachevalier & Loveland, 2006; Bast et al., 2018; Dichter et al., 2012; Huang et al., 2021; Kim et al., 2022; Polzer et al., 2022; Supekar et al., 2018). Differences in the orbitofrontal-amygdala circuit, a circuit integral to motivation, have been found to be connected to the etiology of ASD (Bachevalier & Loveland, 2006). Maturing slightly later than the amygdala, the OFC begins to reach its full potential around the second year of life (Happaney et al., 2004), so it has been postulated that the severity of the emotional and social changes as well as their developmental time course may vary (Happaney et al., 2004). An empirical study found that the autistic individuals showed significantly greater activation than TD individuals in response to the facial photographs in only two regions: the left amygdala and orbitofrontal gyrus, and amygdala activation was strongly and positively associated with, and also predicted by, the amount of eye gaze in autistic individuals (Dalton et al., 2005). This suggests that individuals with ASD display a heightened emotional response to faces, which is likely related to their focus on the eyes. Many studies have been conducted in recent years to discover the neural mechanisms of social reward in individuals with ASD, most of them using smiling faces as social reward feedback (Choi et al., 2015; Damiano et al., 2015; Monk et al., 2010; Kohls et al., 2018; Scott-Van Zeeland et al., 2010). Both reduced and increased responses to smiling faces have been reported in reward-related brain regions (Monk et al., 2010) others have not (Choi et al., 2015; Clements et al., 2018; Kohls et al., 2013; Kohls, Thonessen, et al., 2014; Monk et al., 2010).

Another neural systems with alterations related to ASD is the LC-NE system. This system is placed on both sides of the brainstem just beneath the cerebellum and lateral to the fourth ventricle (Poe et al., 2020).The LC-NE system is also closely related to dopamine system (Sara, 2009), which is in turn associated with orbitofrontal-amygdala circuitry (Bachevalier, 2005). A review has proposed that atypical LC-NE activity may be a developmental mechanism leading to reduced social attention as well as social interaction and communication impairments in children with autism (Bast et al., 2018). In terms of markers of LC-NE system activation, many studies have shown that pupil diameter (PD) responses are directly related to LC activity (Alnaes et al., 2014; Gilzenrat et al., 2010; Murphy et al., 2014; Sterpenich et al., 2006). TD children showed an increased PD when looking at happy faces and increased pupil diameter is related to higher emotional arousal and greater rewarding experiences in healthy children when viewing pleasant faces (Sepeta et al., 2012). However, in children with ASD no increased pupil diameter was found when viewing happy faces, leading to a deficit in the capture of reward information and a lack of positive experience (Sepeta et al., 2012). Inconsistent results have also been found with another study reporting no difference between TD and ASD individuals for upright faces although for inverted faces pupil diameter was larger in ASD individuals (Falck-Ytter, 2008). In summary, while many studies demonstrate PD as an explicit index of LC activation (Alnaes et al., 2014; Bast, Boxhoorn, et al., 2023; Bast, Mason, et al., 2023; Keehn et al., 2021; Polzer et al., 2022; Privitera et al., 2020) findings in relation to individuals with ASD in the context of social reward are inconsistent.

Few studies pertaining to the altered neural mechanisms underlying social reward experiences in young ASD individuals with limited language or cognitive abilities have been carried out, with most previous studies focusing on high-functioning adolescents or adults (Clements et al., 2018). Task functional magnetic resonance imaging (fMRI) scanning, which would provide high spatial resolution for researching neural mechanisms, is very challenging in low-functioning individuals with ASD and particularly in toddlers. It is therefore still unclear to what extent altered social reward processing may occur generally in young children with ASD. Functional Near-infrared Spectroscopy (fNIRS) is an alternative rising neuroimaging strategy which may be particularly useful in the context of young and low functioning individuals with ASD due to its portability, relative insensitivity to movement and greater temporal resolution (Ferrari and Quaresima, 2012). In recent years, some studies have used fNIRS technology to explore neural activation in high-risk infants for autism, and it was found that they showed lower activation of some regions of the frontal and temporal cortex than their siblings when viewing or hearing social information versus non-social information (Braukmann et al., 2018; Lloyd-Fox et al., 2018; Pecukonis et al., 2021). However, no studies have yet been conducted on happy faces using this technology in order to investigate the possibility of altered social reward processing in young ASD compared with TD individuals.

### 1.3 The current study

Given this background of previous studies we have therefore investigated differences in eye-gaze and neural mechanisms between young children with ASD relative to TD when viewing happy faces and simplified happy face expressions in the form of emoticons, and if they are associated with adaptive abilities and autistic symptoms. If certain indices were found to be reliable in predicting adaptive difficulties and symptom severity in ASD, it could help to create an efficient and objective approach to predict adaptive functioning and requirement of extra services and support.

We hypothesized that children with ASD would show different gaze patterns to happy faces and eye-mouth viewing patterns compared to TD children particularly when presented with more complex natural facial expressions. We expected that TD children would pay more attention to the whole faces especially eyes, which contain more social information, whereas individuals with ASD would not show this preference in accordance with the eye avoidance and excess mouth/diminished eye gaze hypotheses. On the other hand, when presented with simple emoticon faces, we predicted that the gaze patterns of ASD and TD individuals would be similar, both paying similar attention to the eyes. Furthermore, we anticipated that the differences in viewing patterns between the two conditions would be related to the severity of social symptoms in autistic individuals. In accordance with the social motivation hypothesis, we predicted that individuals with ASD would have a smaller PD and reduced neural activation when presented with happy faces, and the degree of decreased PD and neutral activation would be linked to their social motivation.The Lancet commission has recently suggested a call to action to create more precise labels, like ‘profound autism’, to identify individuals who require extra services and support due to their severe condition (Lord et al., 2022). Prediction of autism based on early snapshots of adaptive functioning should be made more robust objective and prognosis-relevant, thus aiding future research (Mandelli et al., 2023). Thus, we hypothesized that eye-tracking and functional near-infrared spectroscopy (fNIRS) indices could be used to predict adaptive difficulties and severity of social symptoms in young children with ASD.

## 2 Methods

### 2.1 Participants

36 ASD and 36 age-(ASD: M = 48.89 months; SD = 15.36; TD: M = 53.83months; SD = 11.34) and gender-(6 females) matched TD children were recruited in the present study. The study was conducted in the Chengdu Maternal and Children’s Central Hospital (CMCCH) and Sichuan Normal University in accordance with the principles of the Declaration of Helsinki. It was approved by the Ethics Committee of CMCCH as well as the Ethics Committee of the University of Sichuan Normal University (number 202165). Written informed consents were provided by parents or legal guardians of children before participation.

Children diagnosed with ASD were recruited from CMCCH outpatient clinics. Participants were considered eligible if they were diagnosed with ASD according to the Diagnostic and Statistical Manual of Mental Disorders, Fifth Edition (DSM-V)(American Psychiatric Association,2013) by clinicians, aged 2.5-6 years, and their diagnosis was confirmed with the Autism Diagnostic Observation Schedule-2 (ADOS-2 – Lord et al., 2012) administered by research reliable trained individuals. Individuals with genetic disorders, diagnosed with epilepsy, cerebral palsy, attention deficit hyperactivity disorder (ADHD) or other psychiatric disorder were excluded. Children with severe hearing, or visual deficits were not included. TD children were recruited from Experimental Kindergarten of Sichuan Normal University and Qingshen Experimental Kindergarten in Sichuan. fNIRS data from 10 ASD and 5 TD individuals could not be included due to acquisition difficulty or poor data quality. All participants’ eye-tracking data were valid using the threshold of fixation duration for the screen being no less than 10s and with successful 5 point calibration criteria (details see SI).

### 2.2 Clinical Assessments

The diagnosis of ASD in participants was confirmed by the gold-standard ADOS-2 (Lord et al., 2012). ADOS-2 is a semi-structured assessment of autism severity administered by an experienced evaluator who conducts a series of activities with the participant in the presence of the primary caregiver and then rates them based on notes taken during the activities. The total score is derived by summing the Social Affect (SA), Restricted and Repetitive Stereotyped Behaviors (RRB), which can be used to compare with cut-off scores to determine whether the participant with autism, on the autism spectrum or non-autism. The calibrated severity score (CSS) of 1-10 is converted from the raw total scores to ascertain the severity of symptoms. Modules 1 and 2 were mainly used in this study, but a minimal number of participants were tested by Module T. Additionally, a series of other report questionnaires to assess ASD symptoms and adaptive behavior, including the Social Responsiveness Scale, Second Edition (SRS-2)(Constantino & Gruber, 2012), and the Adaptive Behavior Assessment System-Second Edition (ABAS-2)(Oakland & Harrison, 2011) were conducted (See Table 1).

**Table 1.**
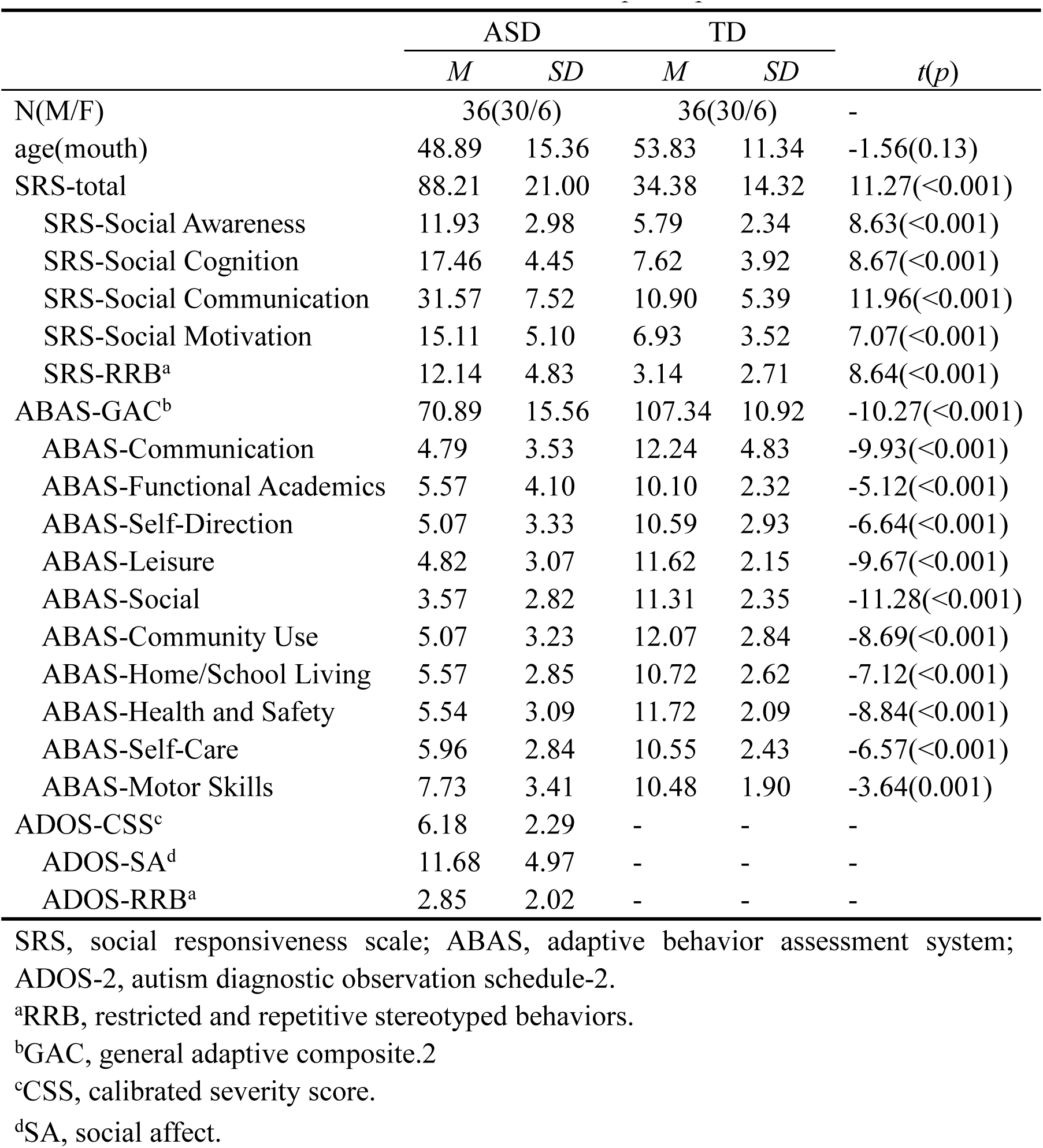
Characteristics of the participants.

|  | ASD |  | TD |  | *t(p)* |
| --- | --- | --- | --- | --- | --- |
|  | *M* | *SD* | *M* | *SD* |  |
| N(M/F) | 36(30/6) |  | 36(30/6) |  | - |
| age(month) | 48.89 | 15.36 | 53.83 | 11.34 | -1.56(0.13) |
| SRS-total | 88.21 | 21.00 | 34.38 | 14.32 | 11.27(<0.001) |
| SRS-Social Awareness | 11.93 | 2.98 | 5.79 | 2.34 | 8.63(<0.001) |
| SRS-Social Cognition | 17.46 | 4.45 | 7.62 | 3.92 | 8.67(<0.001) |
| SRS-Social Communication | 31.57 | 7.52 | 10.90 | 5.39 | 11.96(<0.001) |
| SRS-Social Motivation | 15.11 | 5.10 | 6.93 | 3.52 | 7.07(<0.001) |
| SRS-RRB<sup>a</sup> | 12.14 | 4.83 | 3.14 | 2.71 | 8.64(<0.001) |
| ABAS-GAC<sup>b</sup> | 70.89 | 15.56 | 107.34 | 10.92 | -10.27(<0.001) |
| ABAS-Communication | 4.79 | 3.53 | 12.24 | 4.83 | -9.93(<0.001) |
| ABAS-Functional Academics | 5.57 | 4.10 | 10.10 | 2.32 | -5.12(<0.001) |
| ABAS-Self-Direction | 5.07 | 3.33 | 10.59 | 2.93 | -6.64(<0.001) |
| ABAS-Leisure | 4.82 | 3.07 | 11.62 | 2.15 | -9.67(<0.001) |
| ABAS-Social | 3.57 | 2.82 | 11.31 | 2.35 | -11.28(<0.001) |
| ABAS-Community Use | 5.07 | 3.23 | 12.07 | 2.84 | -8.69(<0.001) |
| ABAS-Home/School Living | 5.57 | 2.85 | 10.72 | 2.62 | -7.12(<0.001) |
| ABAS-Health and Safety | 5.54 | 3.09 | 11.72 | 2.09 | -8.84(<0.001) |
| ABAS-Self-Care | 5.96 | 2.84 | 10.55 | 2.43 | -6.57(<0.001) |
| ABAS-Motor Skills | 7.73 | 3.41 | 10.48 | 1.90 | -3.64(0.001) |
| ADOS-CSS<sup>c</sup> | 6.18 | 2.29 | - | - | - |
| ADOS-SA<sup>d</sup> | 11.68 | 4.97 | - | - | - |
| ADOS-RRB<sup>a</sup> | 2.85 | 2.02 | - | - | - |

SRS, social responsiveness scale; ABAS, adaptive behavior assessment system;
ADOS-2, autism diagnostic observation schedule-2.
<sup>a</sup>RRB, restricted and repetitive stereotyped behaviors.
<sup>b</sup>GAC, general adaptive composite.2
<sup>c</sup>CSS, calibrated severity score.
<sup>d</sup>SA, social affect.

### 2.3 Happy Face Free Viewing Task

To investigate gaze patterns and neural responses, we adapted our previous emoticon versus real face task from Le et al. (2020) into a block design study. The blocks were composed of pictures of a child with a happy child face (HCF) and happy emoticon face (HEF)(See Figure 1). Areas of interest (AOIs) included mouths and eyes of child and emoticon faces (See Figure 1). In this paradigm, stimuli in each block were displayed for 2 seconds, 5 stimuli per block and 4 block per condition. Twenty stimuli were used in the current study. Each block was separated by a jittered interval (an average of 8 seconds).

**Figure 1.**
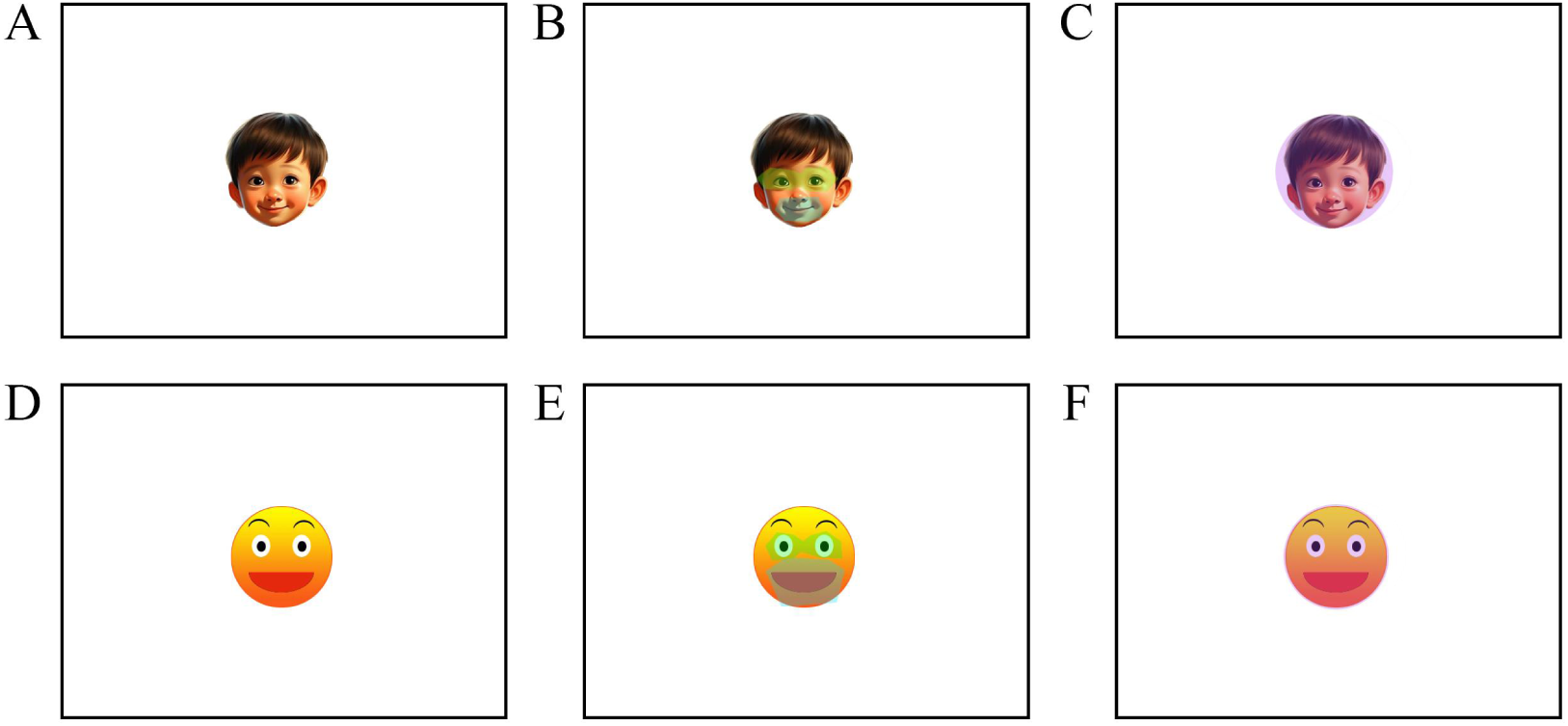
(A) An example in the happy complex face (HCF) condition. Of note, the present facial expression is for illustration only and not included in the original stimulus set. **(B)** A schematic representation of the areas of interest (AOIs) for the eyes and mouth in the HCF condition is shown; **(C)** The AOI of the whole face in the HCF condition; **(D)** An example in the happy emoticon face (HEF) condition; **(E)**The AOIs of the eyes and mouth delimited in the HEF condition; **(F)** The AOI of the whole face in the HEF condition.

### 2.4 Data acquisition

#### Eye-tracking data

The children were seated by themselves in front of a monitor, with no instructions from the caregivers or investigator, so that they could observe the images on the screen at their own pace. Eye-tracking data was collected using a Tobii Pro Spectrum screen-based remote eye tracker as the hardware device with Tobii Pro Lab (version 1.142.27188) as the software to record eye gaze patterns, with a sampling rate of 120Hz. E-prime 3.0 linked with the Tobii software was used for stimulus presentation. A standard five-point calibration was applied to conduct the eye-tracking calibration (See SI). If the calibration quality did not meet the required standard, children had to keep repeating the process until they passed. The average of binocular eyes was then used to determine the fixation parameters. The eye-tracking monitor, a TFT-LCD with a resolution of 2300, 1920×1080, had a refresh rate of 60 Hz and a brightness of 100%.

#### fNIRS data

While children viewed stimuli in the current study, we used Nirsit-Lite (Kids), a portable functional Near-Infrared Spectroscopy (fNIRS), to collect the hemodynamic changes in the OFC (OBELAB Inc., Seoul, Republic of Korea)(Bak, Shin, et al., 2022;Cho et al., 2022; Choi et al., 2016; Jang et al., 2020). The device has five dual-wavelength (780 and 850 nm) laser diodes and seven photodetectors comprising 15 channels, which is designed according to the standard International 10-20 EEG system and the optode distance is 2.5cm with 8.138Hz sampling rate (Bak, Shin, et al., 2022; Cho et al., 2022; Choi et al., 2016; Jang et al., 2020). The alignment of channels is shown in Fig. 3A.

### 2.5 Data analysis

#### Eye-tracking data analysis

We plotted the areas of interests (AOIs) in the Tobii Pro Lab for the HCF condition and the HEF condition, respectively, and schematic diagrams are presented in Figure 1. In each condition, an eye area, a mouth area, and an entire face area were drawn, and the entire face area was an identical size for the two conditions. To ensure that the fixation at the edge of the region was accurately measured, the plotted AOI was slightly larger than the corresponding actual region. Fixation data was exported from Tobii Pro Lab using the I-VT fixation filter (Details see SI).

The following metrics were utilized: gaze pattern to child and emoticon faces was quantified as the mean proportion of total fixation time or fixation counts spent on looking at whole faces and the eye and mouth region as in previous face processing tasks (Kwon et al., 2019; Le et al., 2020). We measured mean proportion of fixation counts and mean individual fixation duration. Pupil diameter (PD) in each condition was measured to quantify children’s arousal level in response to the faces. The proportion of the total fixation time was calculated as follows: % total fixation time spent on faces (TFD%(W)) = (time spent on faces/total looking time on stimuli screens)×100; % total fixation time on eye-mouth gaze pattern (TFD%(E-M)) to child faces = (time spent on eyes-mouths/total looking time on stimulus screens). The FC%(W) and FC%(E-M) indices were calculated accordingly. Two-way ANOVA tests were conducted to investigate differences, followed by Bonferroni-corrected post hoc comparisons. Additionally, main effects between groups were also assessed.

#### fNIRS data analysis

The NIRS-KIT (Hou et al., 2021), a MATLAB toolbox, was used for fNIRS signal processing. Four steps were applied were applied for pre-processing in accordance with a previous study (Hou et al., 2021): 1) trimming the signals during the non-task time to leave the actual task interval; 2) linear-detrending method; 3) correlation-based signal improvement (CBSI) was used for head motion correction; 4) infinite impulse response (IIR) filter was applied for band-pass with cutoff frequencies at 0.01 Hz and 0.10 Hz. Subsequently, the general linear model (GLM) was employed to characterize the hemodynamic response of oxygenated hemoglobin (HbO) under the two conditions. HbO was chosen as the index to be analyzed since its fluctuations are more sensitive than deoxygenated hemoglobin (HbR).

Similarly, ANOVA tests were carried out for the HbO value of fNIRS data, followed by post hoc tests for channels exhibiting significant interactions and main effects between groups. Bonferroni correction of HbO values from 15 channels to control alpha error was calculated (*p_Bonferroni_* < 0.05/15 was considered significant for the interaction and main effects).

#### Correlation between group differences and adaptive difficulties/ autistic symptoms

To investigate the relationship between the distinct neural and behavioral patterns and participants autistic symptoms and adaptive abilities, Pearson correlations were calculated between distinct eye-tracking and fNIRS variable patterns and scores of SRS-2, ABAS-2 and ADOS-2. For the purpose of exploring whether the distinct eye-tracking and fNIRS indexes in this study were useful predictor of autism-related severity symptoms and adaptive behaviors in children, multiple linear regression analyses were performed on the total score or social-relevant dimensions of the SRS-2 and ABAS-2 using metrics from eye-tracking and fNIRS as independent variables to verify their potential categorical or predictive utility.

#### Classification markers for ASD

A Support Vector Machine (SVM)-based machine learning classification test was conducted to investigate whether the eye-tracking and fNIRS data from the present task could be used as valid biomarkers to detect ASD. We scheduled to select all the statistical values with between-group discrepancies as classification metrics. Linear-kernel SVMs have been employed for group classification. The SVMs have been trained according to the leave-one-out cross validation (LOOCV) technique; hence, excluding one participant from the training set at each iteration, and testing the trained SVM on it.

In addition to the classification accuracy of the SVM (percentage of correctly classified participants), we also evaluated the receiver operating characteristic (ROC) curves for classification based on each significant variable, where the sensitivity% (i.e., percentage of individuals diagnosed with ASD patients were correctly classified as having ASD) and the 100%-specificity% (i.e., percentage of incorrectly classified TD control subjects) were plotted as the horizontal and vertical coordinates, respectively. From the ROC curves, the area under the curve (AUC) was calculated to indicate the classification efficiency.

## 3 Results

### 3.1 Group differences in gaze patterns

No significant main or interaction effects (*ps* > 0.668) were found for both TFD%(W) and FC%(W). However, significant interactions were found for eye-mouth gaze patterns [TFD%(E-M) (F_1,70_ = 5.237, *p* = 0.025), FC% (E-M) (F_1,70_ = 6.362, *p* = 0.014)]. Post hoc analysis showed that within the TD group there was a significant difference [TFD(E-M)%: t_35_ = –2.396, *p* = 0.022), FC(E-M)%: t_35_ = –2.136, *p* = 0.040] with children focusing more on the eyes in the HCF condition and more on the mouth in the HEF one. However within the ASD group, there was no significant difference (*ps* > 0.170). The ANOVA analysis of PD revealed significant main effects of group (F_1,70_ = 3.237, *p* =0.050) and stimuli (F_1,70_ = 6.280, *p* < 0.001), and a significant interaction (F_1,70_ = 13.059, *p* = 0.001), with post hoc tests showing that PD was larger in the ASD than TD group in the HCF condition (*p* = 0.013) but not in the HEF condition (*p* = 0.200). Children in both groups had larger PD looking at HCF than HEF [ASD (*p* < 0.001), TD (*p* = 0.025)]. This indicates that the ASD group showed a significant greater PD than the TD group when gazing at HCF, while the two groups did not show a significant difference in PD when gazing at HEF (See Figure 2).

**Figure 2.**
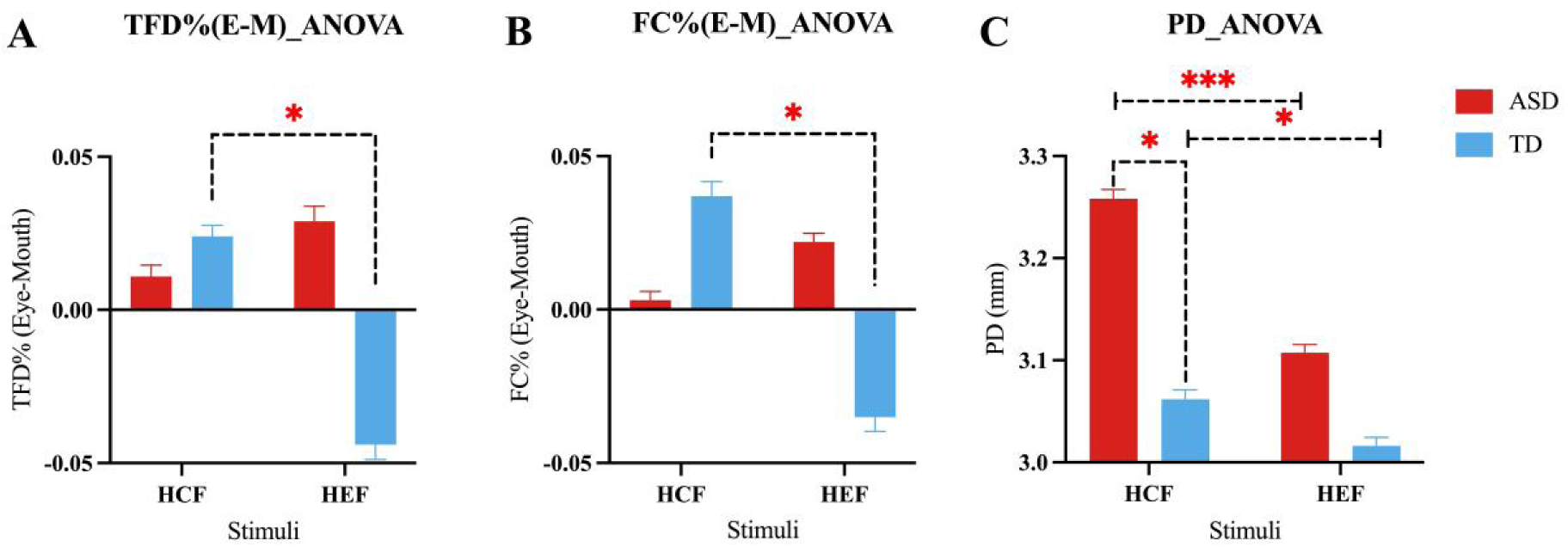
ANOVA results of fixation patterns and pupil diamter (PD). **(A)** TFD (total fixation duration) % (E-M) ANOVA result; **(B)** FC (fixation count) % (E-M) ANOVA result; **(C)** PD ANOVA result. * *p*<.05, \*\*\**p*<0.001

**Figure 3.**
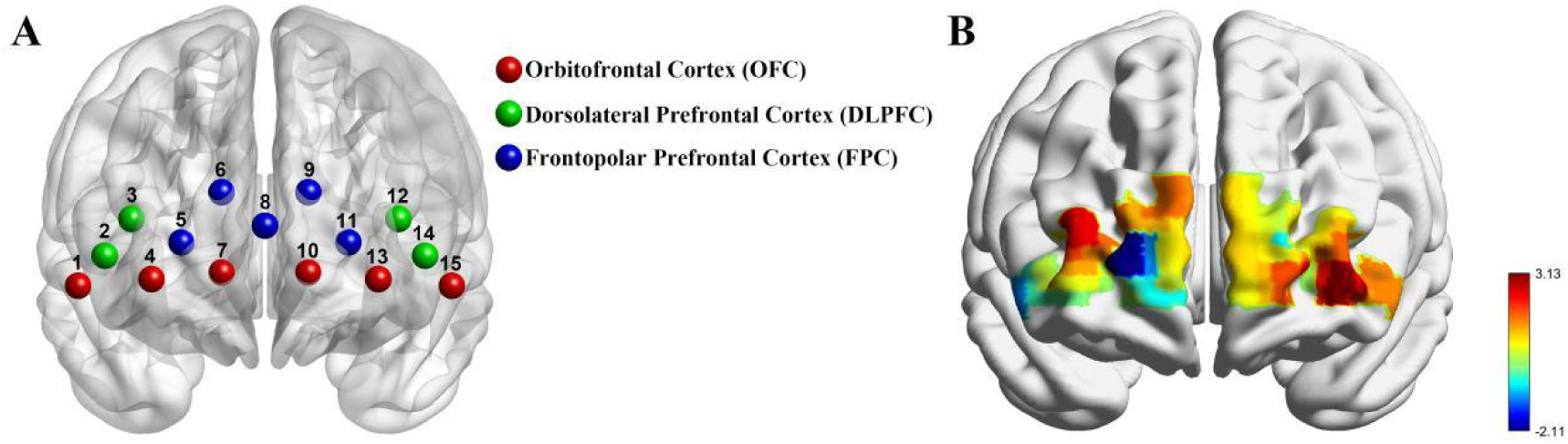
**(A)** The alignment of OFC channels is shown. **(B)** Schematic diagram showing a pseudocolor representation of the oxy-HbO concentration per channel for ASD-TD averaged over HCF and HEF conditions.

### 3.2 Group differences of neural responses

ANOVAs revealed no significant interactions on any of the channels after multiple comparison correction by Bonferroni correction (*ps* > 0.024), but there was a significant main effect between groups. As shown in Figure 3, on channels 13, there was a significant difference in HbO concentration between the two groups (t_56_ = 3.126, *p* = 0.0028), indicating an increased OFC response in the ASD group compared to the TD group (See Figure 3).

### 3.3 Correlation

The results presented above for the analysis of eye-tracking and fNIRS disclosed that there were significant differences in eye-mouth gaze pattern, PD and OFC activation (channel 13 of fNIRS) between the ASD and TD groups. Firstly, their correlation with SRS-2 was examined and results showed both eye-mouth gaze pattern, pupil diameter measures and mean HbO values of fNIRS channels were significantly correlated with the total SRS-2 score. As shown in Figures 2 and 4 (HCF-HEF) TFD%(E-M) (r = –0.328, *p* = 0.013), (HCF-HEF)FC%(E-M) (r = –0.383, *p* = 0.003) were negatively correlated with SRS-2 total score. (HCF-HEF)PD (r = 0.330, *p* = 0.012) and OFC activation of channel 13 (r = 0.467, *p* = 0.001) was positively correlated with SRS-2 total score. The detailed correlation analysis between eye-tracking, OFC activation and each dimension of SRS-2 is shown in Table S1.

**Figure 4.**
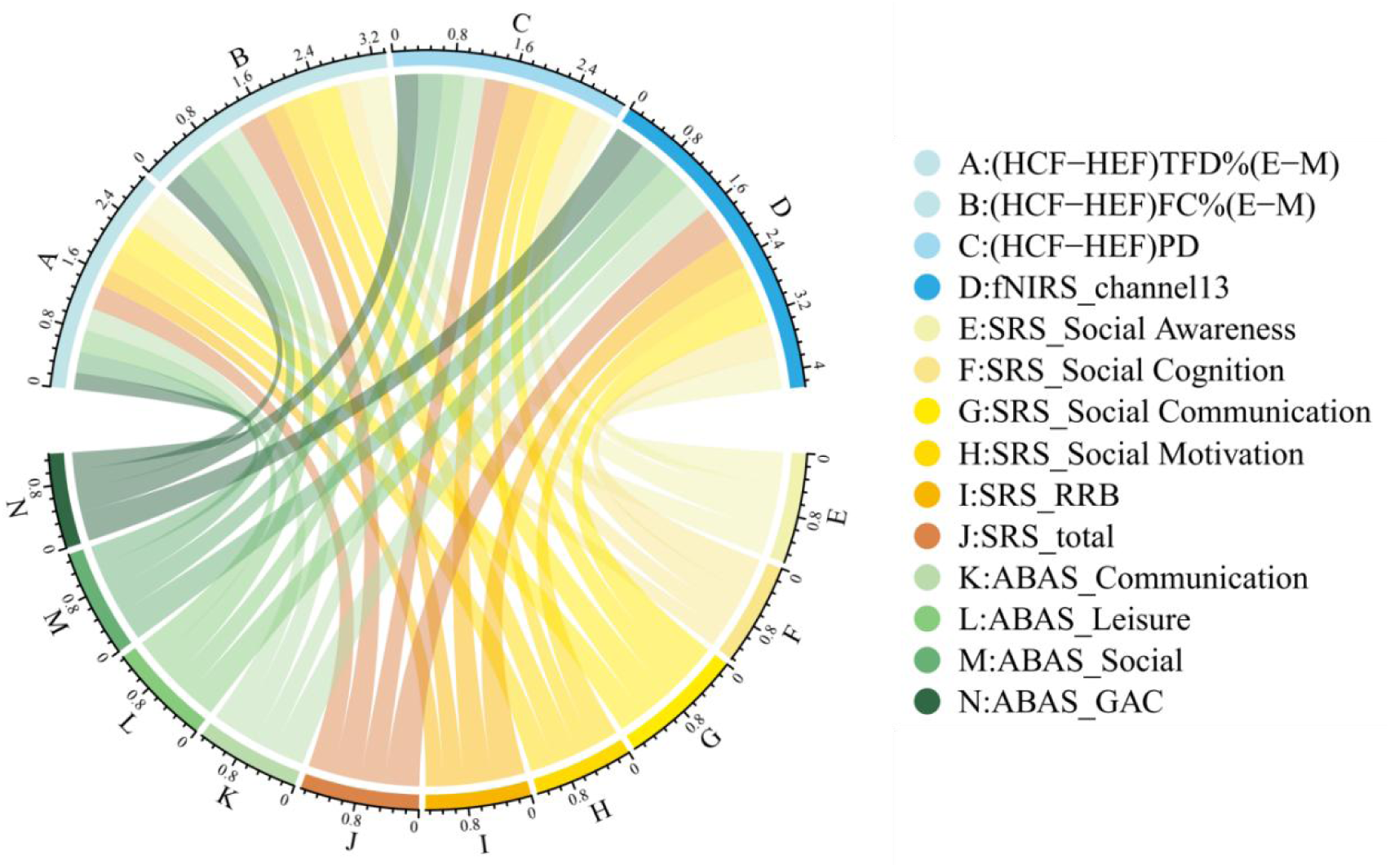
The results of the correlation analysis between eye-tracking, channel 13 fNIRS data and SRS-2, parts of ABAS-2 scores. For ADOS-2, the results showed that PD scores were significantly correlated with the social affect dimensions [HCF_PD (r = 0.368, *p* = 0.032), HEF_PD (r = 0.404, *p* = 0.018), mean_PD (r = 0.390, *p* = 0.023)] and CSS [HCF_PD (r = 0.338, *p* = 0.051), HEF_PD (r = 0.367, *p* = 0.033), mean_PD (r = 0.335, *p* = 0.039)] (except for CF_PD which nearly achieved significance with ADOS_CSS). Details see Table S3.

Eye-tracking and fNIRS data were then examined for correlations with ABAS-2. As shown in Figure 4 and Figure S2, Eye-mouth gaze patterns were found to be significantly correlated with children’s social skills, including leisure [TFD%(E-M) (r = 0.299, *p* = 0.024), FC% (E-M) (r = 0.334, *p* = 0.011)] and social [TFD%(E-M) (r = 0.279, *p* = 0.036), FC% (E-M) (r = 0.353, *p* = 0.007)], as well as the “communication” dimension [TFD%(E-M) (r = 0.299, *p* = 0.024), FC% (E-M) (r = 0.320, *p* = 0.015)]. Pupil diameter difference scores of HCF versus HEF were negatively correlated with most dimensions of adaptive skills (rs < –0.277, *ps* < 0.038) as well as with total scores (r = –0.313, *p* = 0.018). Neural response data was negatively correlated with almost all dimensions (rs < –0.296, *ps* < 0.042) of ABAS-2 as well as with total scores (r = –0.402, *p* = 0.005). For details see Table S2.

### 3.4 Classification markers

Machine learning analysis was performed on gaze patterns, PD and the mean HbO values of fNIRS channels 13 which differed significantly between the two groups and led to a classification accuracy of 66.67%. For the ROC analysis fNIRS channel 13 proved to be the most effective in achieving an AUC of 74.3% (95% CI. 61.3%-87.3%), sensitivity of 73.1% (95% CI, 53.9%-86.3%) and specificity of 67.7% (95% CI, 50.1%-81.4%). The other three variables yielded AUCs of 58.3%, 61.9%, 71.9%, respectively. (see Figure 5)

**Figure 5.**
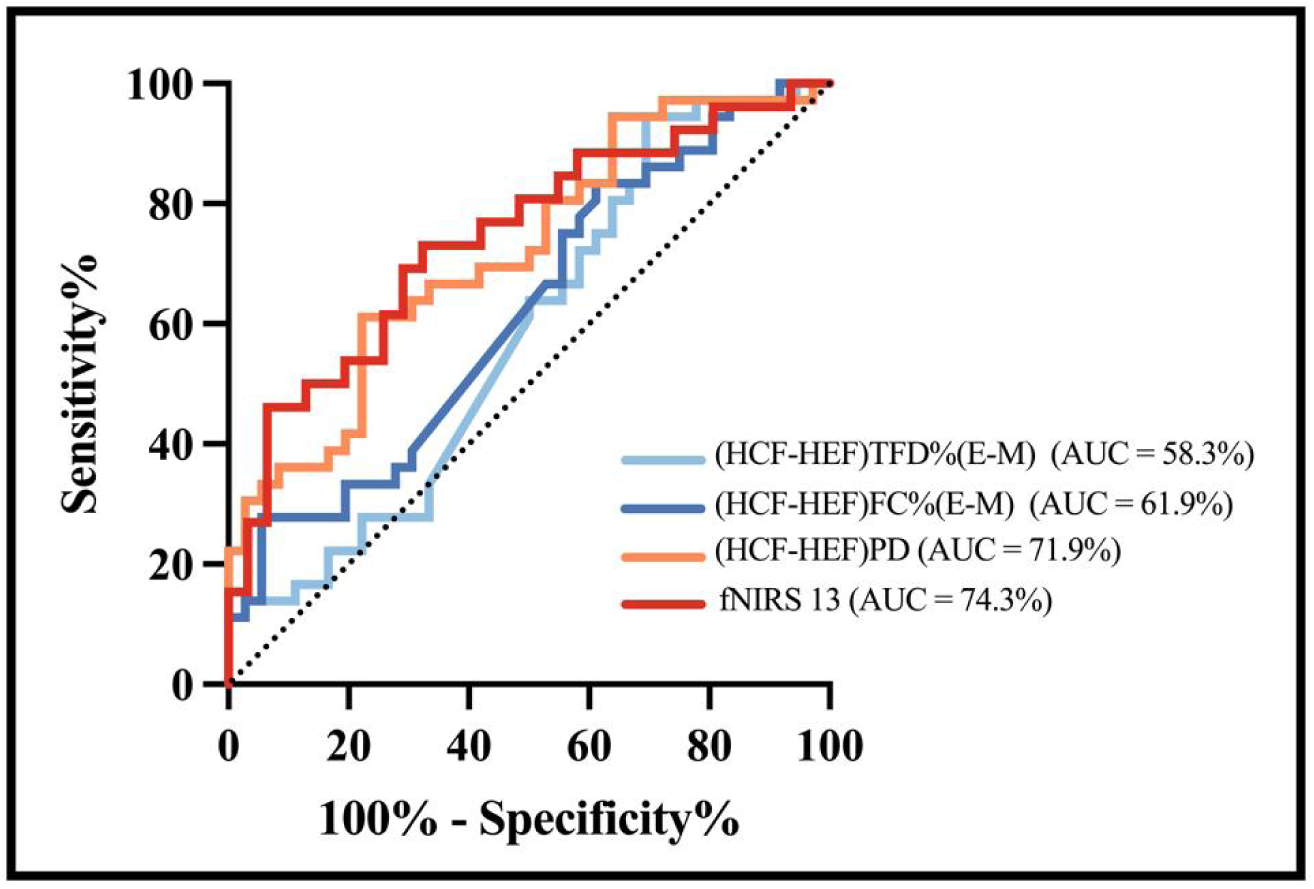
ROC curve plot graphically illustrating the true-positive rate (sensitivity%) vs. the false-positive rate (100% – specificity%)

A similar analysis of clinical symptom severity scores ADOS-2 in ASD group showed that pupil diameter scores were significantly correlated with the social affect dimensions [HCF_PD (r = 0.368, *p* = 0.032), HEF_PD (r = 0.404, *p* = 0.018), mean_PD (r = 0.390, *p* = 0.023)] and symptom comparison CSS scores [HCF_PD (r = 0.338, *p* = 0.051), HEF_PD (r = 0.367, *p* = 0.033), mean_PD (r = 0.335, *p* = 0.039)] (except for CF_PD which nearly achieved significance with ADOS_CSS). Details see Table S3.

### 3.5 Predictors of Autistic Symptom and Adaptive Ability of Children

The results of the above correlation analysis demonstrated that the total score of SRS-2 was significantly correlated with the eye-mouth gaze pattern, pupil diameters, and neural response of the OFC. To investigate whether they can predict the total score of SRS-2 multiple linear regression was conducted. To avoid the possible effects of multi-collinearity, (HCF-HEF)FC%(E-M) was selected as a representative of the fixation pattern, the neural response from channel 13 of fNIRS was selected as representative of fNIRS data and for HCF-HEF for PD. These three were used as independent variables and the total score of SRS-2 as the dependent variable. The results showed that all these three had a good predictive effect with an R-squared of 0.405, which means that the predictive rate of the three on the total SRS-2 achieved 40.5% (See Table 2).

**Table 2.** Multiple Linear Regression on SRS-2 Results.

| | *b* | *SE* | $\beta$ |
| --- | --- | --- | --- |
| (CF-EF)FC%(E-M) | -84.36 | 31.63 | **-0.314\*** |
| CF-EF_PD | 67.30 | 28.37 | **0.284\*** |
| Channel 13 | 4260798.90 | 1406401.79 | **0.363\*\*** |
| $\Delta R^2$ | | **0.405\*\*\*** | |

Note: \* $p < .05$ , \*\* $p < .01$ , \*\*\* $p < .001$

Analogous analyses were applied to the three ABAS-2 dimensions significantly associated with sociability (communication, leisure, and social) as well as ABAS total scores. The results showed that (HCF-HEF)FC%(E-M), (HCF-HEF)_PD and channel 13 were significant predictors of the social dimension of the ABAS-2, with an R-squared of 0.336. The specific values are shown in Table 3. (HCF-HEF)FC%(E-M) and channel 13 were also significant predictors of the ABAS total scores, with an R-square of 0.254, as shown in Table S4.

**Table 3.** Multiple Linear Regression on ABAS-Social.

| | b | SE | $\beta$ |
| --- | --- | --- | --- |
| (CF-EF)FC%(E-M) | 11.587 | 4.669 | **0.309\*** |
| CF-EF_PD | -8.550 | 4.188 | **-0.258\*** |
| Channel 13 | -509367.44 | 207585.945 | **-0.311\*** |
| $\Delta R^2$ | | **0.336\*\*\*** | |

Note: \* $p < .05$ , \*\* $p < .01$ , \*\*\* $p < .001$

## 4 Discussion

The main purpose of the current study was to investigate differences in gaze, patterns, PD and OFC activation during presentation of rewarding happy faces between young children diagnosed with ASD compared with TD children. Importantly, analyses of differences in these measures and their association with distinct domains of adaptive difficulties and autistic symptoms were also performed. Overall, our results show that although there were no significant group differences in time spent viewing the happy faces or the total number of fixations on them, children with ASD showed a greater PD and OFC activation across happy face and emoticon conditions indicating higher arousal and reward valuation of happy faces. TD children showed a preference for more informative rewarding facial features (such as the eyes relative to mouths of real faces and mouths relative to eyes for emoticon faces), whereas children with ASD did not. Gaze patterns towards the eyes and mouth, PD and OFC activation could be used as effective predictors for social adaptive difficulties (R^2^ = 0.336) and the severity of autistic symptoms (R^2^ = 0.405). Higher adaptive abilities and lower autistic symptom scores were predicted by greater attention to informative reward facial features and with reduced PD and OFC responses. Finally, OFC activation when viewing happy faces can be used as a potential biomarker to differentiate ASD and TD groups (with the largest effect sizes and areas under the ROC curve—0.74). Our results fill an important knowledge gap regarding atypical reward face perception processing in young children with ASD, which is inconsistent with the social motivation hypothesis.

### Increased behavioral and neural responses to happy faces in ASD

The greater PD is observed in both TD and ASD groups in the current study provides evidence for hyper-arousal to happy faces per se which is consistent with previous studies to some extent but inconsistent with the social motivation hypothesis. Wagner et al. (2016) reported that infants more likely to be diagnosed subsequently with ASD exhibited greater PD to emotional faces than infants in a low risk group and Reisinger et al. (2020) found that individuals in an ASD group had an increased pupil reactivity compared to those in a TD group. However, a meta-analysis of the pupil response in individuals with ASD only showed a longer latency of the pupil response in ASD groups whereas evidence on other pupil size related indices was conflicting (de Vries et al., 2021). A qualitative evaluation using the framework method suggests that the group differences between ASD and TD could be due to the involvement of the autonomic nervous system and more specifically the locus coeruleus-norepinephrine system (de Vries et al., 2021). Neurons of the brainstem nucleus LC are the sole source of NE (Sara, 2009) and the release of NE throughout the mammalian brain is considered important for modulating attention, arousal, and cognition during many behaviors (Schwarz & Luo, 2015; Usher et al., 1999). The LC-NE system is also closely related to dopamine system, which is closely associated with the orbitofrontal-amygdala circuit (Bachevalier, 2005). For the relation between the LC-NE system and dopamine, Sara (2009) concluded that there were striking similarities between the factors that govern the activity of dopaminergic and NE neurons, suggesting that dopamine and NE were released simultaneously. Additionally, a recent resting-state fMRI study found that ASD individuals had reduced functional connectivity between the NE and dopamine systems (Huang et al., 2021). All above suggest that the LC-NE system may be integrally related to reward processing and one of the key neural differences in ASD.

In addition, results of the current study indicate that autistic children displayed hyper OFC activity when looking at happy faces and emoticons, corroborating earlier research that reported hyper-amygdala activity when ASD individuals viewed fearful, neutral, or ambiguous expressions (Stuart et al., 2022). However, in comparison to their TD counterparts, a magnetoencephalography (MEG) study found that adolescents in the ASD group, exhibited significantly stronger cerebellar activation in response to happy faces (Styliadis et al., 2021). Activation of the amygdala and amygdala-prefrontal connectivity during presentation of happy emotional faces have also been studied in ASD. Monk et al. (2010) presented picture pairs (happy-neutral, sad-neutral, angry-neutral and neutral-neutral expressions) to participants and results indicated that for happy-neutral faces, the ASD group showed greater right amygdala activation relative to the TD control group, and activation in the ASD group was greater for happy-neutral face pairs relative to neutral-neutral face pairs. There was also a positive association with right amygdala activation to the happy-neutral face expression pairs compared to neutral-neutral expression face pairs. For amygdala-ventromedial prefrontal cortex connectivity during the viewing of emotional faces, participants with ASD showed greater positive connectivity relative to the TD control group in the contrast of happy-neutral face expression pairs versus neutral-neutral face expression pairs (Monk et al., 2010). The presence of such altered activation towards happy faces in individuals with ASD implies that the orbitofrontal-amygdala circuit may be associated with their dysfunctional social reward experience. It therefore seems that individuals with ASD may view all social stimuli (even non-negative ones) as threatening.

### Distinct eye-mouth gaze patterns without altered viewing of whole happy expression faces in autistic children

A previous study suggested that children with ASD took longer to perceive happy faces from the time they were presented to, which may indicate that they may have impaired awareness and perception of social rewards (Sato et al., 2017).

While TD children preferred viewing the eyes (informative emotion regions) to mouths, and emoticon mouths (informative emotion regions) instead of eyes for happy faces, children with ASD children showed no differences. Adaptive abilities were predicted to be higher when more informative reward facial features were processed, but this trend was associated with lower autistic symptom severity scores. Our results provide some support for the eye avoidance hypothesis. It is often argued that a decrease in eye contact serves as a contributing factor to the development of ASD, and this symptom is one of the most commonly seen (Klin et al., 2020; Stuart et al., 2023). Eye contact is an important aspect of social communication, as it helps in understanding and interpreting emotions, intentions, and social cues. The differences in eye-mouth gaze patterns suggest that individuals with autism may have difficulties in processing and responding to social cues conveyed through eye contact. Eight of eleven studies supported the eye avoidance hypothesis in a recent meta-analysis suggesting that eye avoidance may be used to reduce hyper-arousal in autistic individuals (Stuart et al., 2023).

Previous research regarding the gaze pattern in relation to emoticons or cartoons specifically in individuals with autism is limited. Emoticons are simplified human faces, but they carry less cognitive and social information than human faces and have hyperbolic features (Fischer & Herbert, 2021). Our current results indicated that TD children preferred to view the face features in emoticons with more social emotional information (eg: emoticon mouths rather than eyes). Silva et al. (2015) discovered that autistic children had a preference for cartoon faces over human faces when presented with an emotion recognition task, using an approach/avoidance paradigm. According to a review, studies have shown that individuals with higher levels of autistic traits tend to display a greater affinity for non-human social agents, such as robots, or cartoons, than those with typical human agents (Atherton & Cross, 2018). Men had a marked advantage in recognizing emoticon or emojis, especially those that were negative, while women were superior at recognizing human facial expressions (Dalle Nogare et al., 2023).

Overall therefore or current results tend to support the eye avoidance hypothesis and stress the importance of increased affective arousal but not the social motivation hypothesis in ASD. This means that instead of a lack of motivation, some autistic children could be steering away from social stimuli so as to lessen their tension and discomfort (Yi et al., 2022). Consistent with our observation of increased responsivity to social reward being associated with greater adaptive behavior difficulties and symptom severity in children with ASD a review has shown a link between increased focus on the face/head and eye regions and improved social performance and reduced autism symptom severity (Riddiford et al., 2022). On the other hand, the amount of gaze allocated to the mouth was reliant on the social and emotional components of the scene and the cognitive profile of the participants. This review also supports the use of gaze variables as potential biomarkers of ASD.

### Conclusion and limitations

Our results concluded atypical hyper reward face perception processing without reward face gaze time inconsistent with the social motivation hypothesis in young children with ASD, which needs to be further evaluated in future investigations. Reduced PD and OFC responses to happy faces, but longer gaze time to eyes of happy expression real faces and mouths of happy emoticons emoticon may predict less autistic symptoms and better adaptive abilities.

There are several limitations in the current study. Firstly, static images of faces were presented that may not be analogous to dynamic faces which are frequently used in other research investigating the impact of social interaction. Secondly, to further investigate the neural and arousal responses of individuals with ASD to social reward, additional components such as gestures, vocalizations, and positive social touch should be included. Thirdly, further research is needed to investigate how individuals with autism perceive and understand emoticons or cartoons specifically and how this may influence their eye-mouth gaze patterns. Fourthly, there were only six females in the current study, so generalizations should be made cautiously to the whole ASD population. Finally, other deeper reward-related brain regions such as the nucleus accumbens and caudate could not be recorded from using fNIRS and therefore they may differ from the patterns of response we observed in the OFC.

## Availability of data

The data that support the main group different finding figures related anonymized data of this study are openly available via the Open Science Framework (OSF) Repository https://osf.io/sxkhc/?view_only=6eb51e64a9b54fce909b5ff6eacdc668 Our ethics approval does not permit public archiving of individual anonymized raw data. Those who wish to access the raw data should contact the corresponding author. Access will be granted to named individuals who adhere to the ethical procedures governing the reuse of sensitive data. They must complete a formal data sharing agreement to obtain the data.

## Acknowledgement

We are thankful to the participants and their families for their involvement in our study. We thank the anonymous reviewers for their constructive comments on the previous versions of the manuscript. Author Contributions Mengyuan Yang: Formal analysis, Data Collection, Writing-Original Draft, Writing-Review & Editing, Visualization; Keith M. Kendrick:Writing-Review; Lihui Huang: Resources; Hong Li & Yi Lei: Resources, Funding acquisition; Lan Zhang: Data collection, Resources; Zijie Wei:Data collection; Pingping Zhang: Data collection; Juan Kou: Conceptualization, Data collection, Methodology, Resources, Writing-Review, Funding acquisition, Supervision.

