## Supplemental file for "Altered Orbitofrontal Cortex Activation and Gaze Patterns to Happy Faces in Autistic Children Predict Adaptive Difficulties but Challenge the Social Motivation Hypothesis"

**Supplementary information**

### Supplementary Methods

#### Calibration process for eye-tracker

The Tobii two-point calibration program was used as in previous studies [Pierce et al., 2015; Kou et al., 2019] for the calibration procedure. We changed the target dots in the program into animated graphics with sound to help attract the children's attention. Calibration was considered to be successful when both eyes had good mapping on all five test positions (See Fig.S1 below). When the calibration accuracy at one or more target locations was greater than 0.5 degrees, the calibration procedure was repeated at those target locations similar to previous studies [Pierce et al., 2015; Kou et al., 2019].

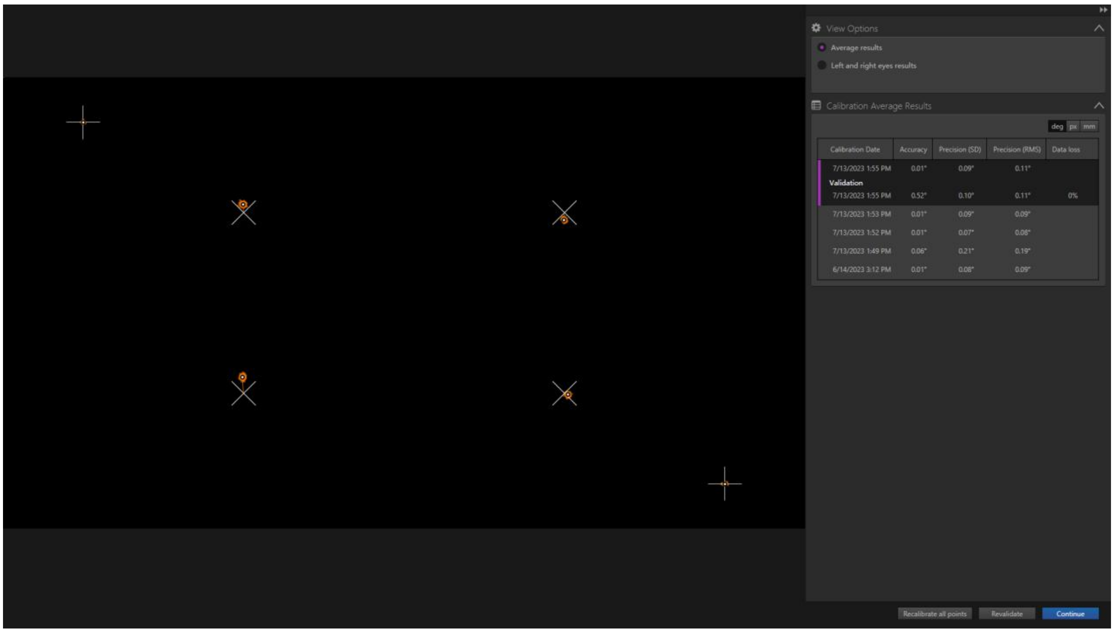


Fig.S1 Successful calibration map sample

#### Details for the parameters of the I-VT filter

Fixations were defined based on the I-VT fixation filter. The parameter settings were as follows: missing gaze data were filled in using linear interpolation, with a maximum gap length of 75ms. Average gaze positions of the left and the right eyes were used to calculate fixations. Noise reduction was moving median and the windows size was 3 samples. The velocity calculator was set at 20 ms. The velocity threshold was set at 30°/s. Adjacent fixations were merged, with the maximum time between merged fixations set to 75 ms and the maximum angle between merged fixations set to 0.5°. Finally, fixations shorter than 60 ms were discarded. [Olsen 2012].

### Supplementary Results

#### Group differences of each AOI viewing

No significant interactions [TFD%(W) ($F_{1,70}$ = 0.002, $p$ = 0.960), FC% (W) ($F_{1,70}$ = 0.065, $p$ = 0.800)] nor main effects [TFD%(W) ($t_{70}$ = = -0.431, $p$ = 0.668), FC% (W) ($t_{70}$ = -0.381, $p$ = 0.704)] were found for both TFD%(W) and FC%(W). No significant interactions [TFD%(E) ($F_{1,70}$ = 2.156, $p$ = 0.147), FC% (E) ($F_{1,70}$ = 2.325, $p$ = 0.132)] nor main effects [TFD%(E) ($t_{70}$ = -0.264, $p$ = 0.793), FC% (E) ($t_{70}$ = -0.057, $p$ = 0.955)] were found for both TFD%(E) and FC%(E), neither. However, significant interactions were found for FC%(M) ($F_{1,70}$ = 4.798, $p$ = 0.032). Post hoc tests for FC%(M) showed that within the TD group, there was a significant difference ($p$ < 0.001) as to HCF and HEF, indicating that the TD children focused more on the mouth in the HEF than the HCF condition, while the ASD group did not show this preference (p = 0.113). Though TFD%(M) showed the same tendency as FC%(M), the result of analysis for TFD%(M) didn't find a significant interaction ($F_{1,70}$ = 3.209, $p$ = 0.078) or main effect ($t_{70}$ = = -0.772, $p$ = 0.443).

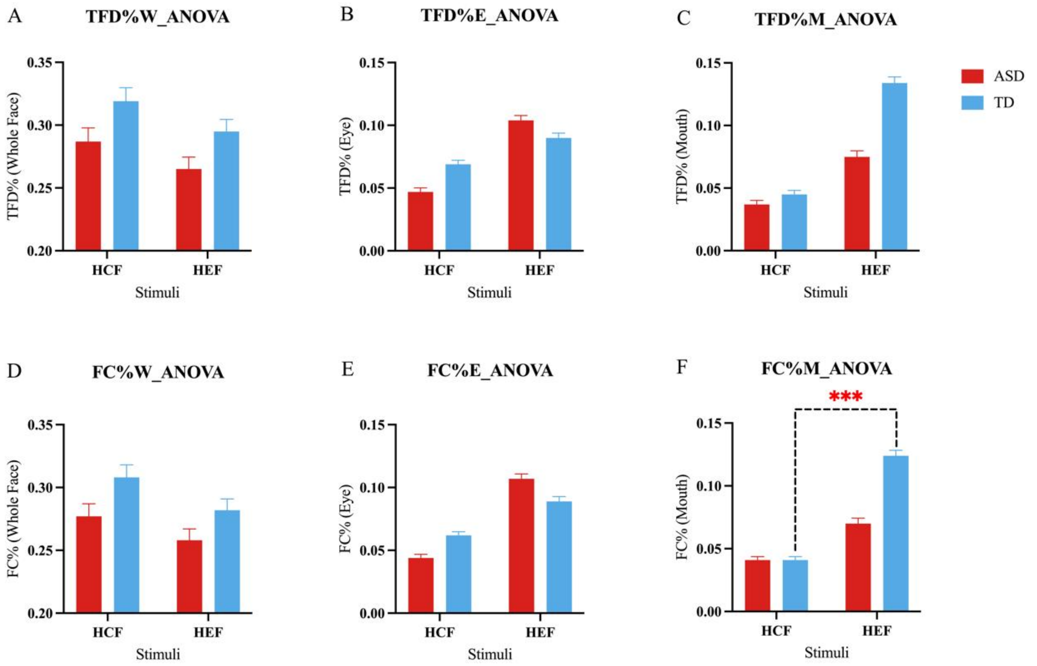


Fig.S2 The ANOVA results of each AOI viewing time and counts. (A, D: for whole face; B, E: for eyes; C, F: for mouth)

**Heatmap**

As supplemental material to Figure 5, the following is a heatmap of the correlation of each of the significant metrics between group of eye-tracking, fNIRS data with all dimensions of the SRS-2 and the socially relevant dimensions and the GAC of the ABAS-2.

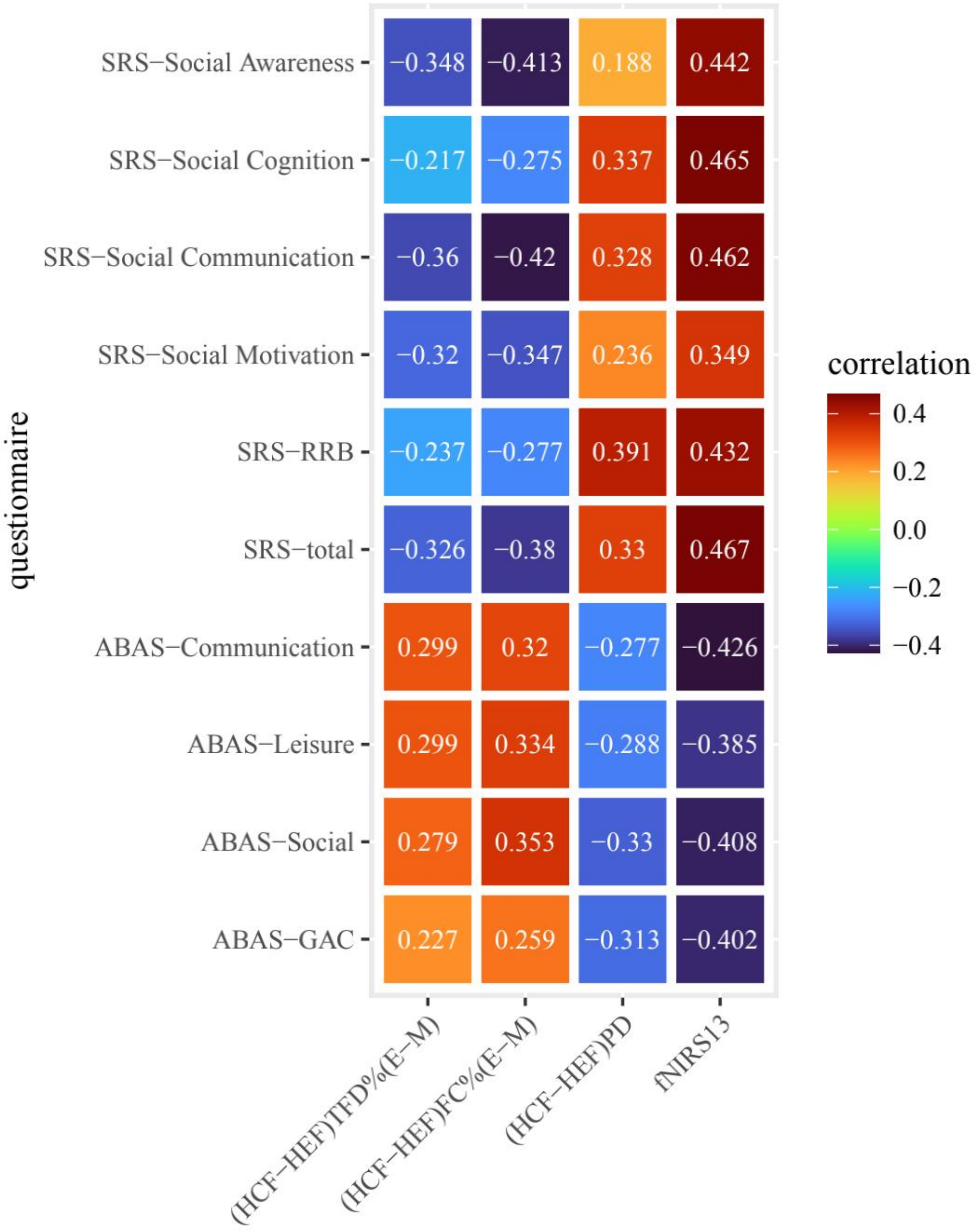


Fig.S3 Heatmap

Table S1

Correlation between eye-tracking, fNIRS data with SRS-2 scores

| | 1 | 2 | 3 | 4 | 5 | 6 | 7 | 9 | 10 | 11 | 12 | 13 | 14 |
|---|---|---|---|---|---|---|---|---|---|---|---|---|---|
| (HCF-HEF)TFD%(E-M) | 1 | | | | | | | | | | | | |
| (HCF-HEF)FC%(E-M) | 0.945$^{***}$ | 1 | | | | | | | | | | | |
| (HCF-HEF)PD | 0.170 | 0.127 | 1 | | | | | | | | | | |
| HCF_PD | 0.125 | 0.052 | 0.455$^{***}$ | 1 | | | | | | | | | |
| HEF_PD | 0.065 | 0.002 | 0.067 | 0.919$^{***}$ | 1 | | | | | | | | |
| mean_PD | 0.098 | 0.029 | 0.277$^{*}$ | 0.982$^{***}$ | 0.977$^{***}$ | 1 | | | | | | | |
| fNIRS_channel13 | -0.160 | -0.199 | 0.163 | 0.057 | -0.009 | -0.026 | 1 | | | | | | |
| SRS-Social Awareness | -0.348$^{**}$ | -0.413$^{**}$ | 0.188 | 0.155 | 0.079 | 0.122 | 0.442$^{**}$ | 1 | | | | | |
| SRS-Social Cognition | -0.217 | -0.275$^{*}$ | 0.337$^{*}$ | 0.162 | 0.015 | 0.094 | 0.465$^{***}$ | 0.834$^{***}$ | 1 | | | | |
| SRS-Social Communication | -0.360$^{**}$ | -0.420$^{**}$ | 0.328$^{*}$ | 0.242 | 0.106 | 0.181 | 0.462$^{***}$ | 0.885$^{***}$ | 0.918$^{***}$ | 1 | | | |
| SRS-Social Motivation | -0.320$^{*}$ | -0.347$^{**}$ | 0.236 | 0.212 | 0.118 | 0.171 | 0.349$^{*}$ | 0.770$^{***}$ | 0.792$^{***}$ | 0.858$^{***}$ | 1 | | |
| SRS-RRB | -0.237 | -0.277$^{*}$ | 0.391$^{**}$ | 0.284$^{*}$ | 0.122 | 0.212 | 0.432$^{**}$ | 0.744$^{***}$ | 0.780$^{***}$ | 0.832$^{***}$ | 0.719$^{***}$ | 1 | |
| SRS-total | -0.326$^{*}$ | -0.380$^{**}$ | 0.330$^{*}$ | 0.235 | 0.097 | 0.173 | 0.467$^{***}$ | 0.905$^{***}$ | 0.941$^{***}$ | 0.983$^{***}$ | 0.896$^{***}$ | 0.880$^{***}$ | 1 |

Note：$^{*}p < .05$, $^{**}p < .01$, $^{***}p < .001$

Table $S_2$

Correlation between eye-tracking, fNIRS data with ABAS-2 scores

| | 1 | 2 | 3 | 4 | 5 | 6 | 7 | 9 | 10 | 11 | 12 | 13 | 14 | 15 | 16 | 17 | 18 | 19 |
|---|---|---|---|---|---|---|---|---|---|---|---|---|---|---|---|---|---|---|
| (HCF-HEF)TFD%(E-M) | 1 | | | | | | | | | | | | | | | | | |
| (HCF-HEF)FC%(E-M) | 0.945$^{***}$ | 1 | | | | | | | | | | | | | | | | |
| (HCF-HEF)PD | 0.170 | 0.127 | 1 | | | | | | | | | | | | | | | |
| HCF_PD | 0.125 | 0.052 | 0.455$^{***}$ | 1 | | | | | | | | | | | | | | |
| HEF_PD | 0.065 | 0.002 | 0.067 | 0.919$^{***}$ | 1 | | | | | | | | | | | | | |
| mean_PD | 0.098 | 0.029 | 0.277$^{*}$ | 0.982$^{***}$ | 0.977$^{***}$ | 1 | | | | | | | | | | | | |
| fNIRS_channel13 | -0.160 | -0.199 | 0.163 | 0.057 | -0.009 | -0.026 | 1 | | | | | | | | | | | |
| ABAS-Communication | 0.299$^{*}$ | 0.320$^{*}$ | -0.277$^{*}$ | -0.153 | -0.034 | -0.098 | -0.426$^{**}$ | 1 | | | | | | | | | | |
| ABAS-Functional Academics | 0.201 | 0.166 | -0.119 | 0.015 | 0.072 | 0.044 | -0.228 | 0.806$^{***}$ | 1 | | | | | | | | | |
| ABAS-Self-Direction | 0.180 | 0.212 | -0.393$^{**}$ | -0.073 | 0.107 | 0.014 | -0.353$^{*}$ | 0.680$^{***}$ | 0.607$^{***}$ | 1 | | | | | | | | |
| ABAS-Leisure | 0.299$^{*}$ | 0.334$^{*}$ | -0.288$^{*}$ | -0.148 | -0.023 | -0.090 | -0.385$^{**}$ | 0.861$^{***}$ | 0.741$^{***}$ | 0.798$^{***}$ | 1 | | | | | | | |
| ABAS-Social | 0.279$^{*}$ | 0.353$^{**}$ | -0.330$^{*}$ | -0.227 | -0.088 | -0.164 | -0.408$^{**}$ | 0.822$^{***}$ | 0.737$^{***}$ | 0.820$^{***}$ | 0.891$^{***}$ | 1 | | | | | | |
| ABAS-Community Use | 0.191 | 0.193 | -0.206 | -0.093 | -0.003 | -0.051 | -0.296$^{*}$ | 0.877$^{***}$ | 0.795$^{***}$ | 0.724$^{***}$ | 0.806$^{***}$ | 0.814$^{***}$ | 1 | | | | | |
| ABAS-Home/School Living | 0.140 | 0.166 | -0.293$^{*}$ | -0.083 | 0.049 | -0.020 | -0.328$^{*}$ | 0.740$^{***}$ | 0.689$^{***}$ | 0.798$^{***}$ | 0.809$^{***}$ | 0.801$^{***}$ | 0.815$^{***}$ | 1 | | | | |
| ABAS-Health and Safety | 0.158 | 0.213 | -0.299$^{*}$ | -0.183 | -0.056 | -0.125 | -0.491$^{***}$ | 0.797$^{***}$ | 0.632$^{***}$ | 0.759$^{***}$ | 0.810$^{***}$ | 0.820$^{***}$ | 0.800$^{***}$ | 0.803$^{***}$ | 1 | | | |
| ABAS-Self-Care | 0.138 | 0.142 | -0.277$^{*}$ | -0.165 | -0.047 | -0.111 | -0.333$^{*}$ | 0.679$^{***}$ | 0.724$^{***}$ | 0.653$^{***}$ | 0.664$^{***}$ | 0.715$^{***}$ | 0.724$^{***}$ | 0.775$^{***}$ | 0.739$^{***}$ | 1 | | |
| ABAS-Motor Skills | 0.276$^{*}$ | 0.259 | -0.071 | 0.112 | 0.153 | 0.135 | -0.511$^{***}$ | 0.588$^{***}$ | 0.600$^{***}$ | 0.474$^{***}$ | 0.486$^{***}$ | 0.554$^{***}$ | 0.595$^{***}$ | 0.547$^{***}$ | 0.601$^{***}$ | 0.557$^{***}$ | 1 | |
| ABAS-GAC | 0.227 | 0.259 | -0.313$^{*}$ | -0.126 | 0.012 | -0.061 | -0.402$^{**}$ | 0.901$^{***}$ | 0.840$^{***}$ | 0.853$^{***}$ | 0.910$^{***}$ | 0.924$^{***}$ | 0.919$^{***}$ | 0.898$^{***}$ | 0.891$^{***}$ | 0.831$^{***}$ | 0.677$^{***}$ | 1 |

Note：$^{*}p < .05$, $^{**}p < .01$, $^{***}p < .001$

Table S3

Correlation between eye-tracking, fNIRS data with ADOS-2 scores

| | 1 | 2 | 3 | 4 | 5 | 6 | 7 | 9 | 10 | 11 |
|---|---|---|---|---|---|---|---|---|---|---|
| (HCF-HEF)TFD%(E-M) | 1 | | | | | | | | | |
| (HCF-HEF)FC%(E-M) | 0.945*** | 1 | | | | | | | | |
| (HCF-HEF)PD | 0.170 | 0.127 | 1 | | | | | | | |
| HCF_PD | 0.125 | 0.052 | 0.455*** | 1 | | | | | | |
| HCF_PD | 0.065 | 0.002 | 0.067 | 0.919*** | 1 | | | | | |
| mean_PD | 0.098 | 0.029 | 0.277* | 0.982*** | 0.977*** | 1 | | | | |
| fNIRS_channel13 | -0.160 | -0.199 | 0.163 | 0.057 | -0.009 | -0.026 | 1 | | | |
| ADOS_SA | -0.029 | -0.100 | 0.049 | 0.369* | 0.401* | 0.389* | 0.088 | 1 | | |
| ADOS_RRB | -0.117 | -0.232 | -0.205 | 0.095 | 0.183 | 0.138 | 0.164 | 0.587*** | 1 | |
| ADOS_CSS | 0.059 | -0.067 | 0.058 | 0.339 | 0.364* | 0.355* | 0.049 | 0.892*** | 0.747*** | 1 |

Note：*p < .05, **p < .01, ***p < .001

Table S4

Multiple Linear Regression on ABAS-GAC

| | b | SE | β |
|---|---|---|---|
| (CF-EF)FC%(E-M) | 24.492 | 23.609 | 0.137 |
| CF-EF_PD | -42.282 | 21.176 | -0.267 |
| Channel 13 | -2541285.5 | 1049712.80 | **-0.325**$^{*}$ |
| $\triangle R^2$ | | **0.254**$^{***}$ | |

Note：$^{*}p < .05$, $^{**}p < .01$, $^{***}p < .001$
